# Predicting drug safety and communicating risk: benefits of a Bayesian approach

**DOI:** 10.1101/193169

**Authors:** Stanley E Lazic, Nicholas Edmunds, Christopher E. Pollard

## Abstract

Drug toxicity is a major source of attrition in drug discovery and development. Pharmaceutical companies routinely use preclinical data to predict clinical outcomes and continue to invest in new assays to improve predictions. However, there are many open questions about how to make the best use of available data, combine diverse data, quantify risk, and communicate risk and uncertainty to enable good decisions. The costs of suboptimal decisions are clear: resources are wasted and patients may be put at risk. We argue that Bayesian methods provide answers to all of these problems and use hERG-mediated QT prolongation as a case study. Benefits of Bayesian machine learning models include intuitive probabilistic statements of risk that incorporate all sources of uncertainty, the option to include diverse data and external information, and visualisations that have a clear link between the output from a statistical model and what this means for risk. Furthermore, Bayesian methods are easy to use with modern software, making their adoption for safety screening straightforward. We include R and Python code to encourage the adoption of these methods.

## Introduction

In drug discovery, data-driven decisions are made to progress projects and compounds to clinical development, with the goal of finding the best candidate drug. A screening cascade or funnel is often designed to progressively increase the likelihood of success for a project. Such screening paradigms are used to develop drugs that interact with a target, thereby modulating a disease in a beneficial way. These cascades often start with low-cost high-throughput binding or cellular functional assays to identify structures that associate with the target of interest. Selected compounds then progress into a more complex *in vitro* model to assess their functional activity, followed by *in vivo* studies to show that they display the correct pharmacology and modulate the disease in the intended way. Parallel to this, similar screening cascades will be conducted in drug metabolism and safety to ensure that compounds have the desired pharmacokinetic and safety profiles. To decide which compounds progress through these screens, the predictive and translational value of the assays must be determined. The predictive value could relate to the next assay in the sequence through to the final treatment of a patient.

The decision to progress a new compound or terminate over safety concerns requires data on past compounds and a statistical or machine learning model to turn the data into an actionable prediction. How do we make the best use of available data and communicate risk and uncertainty to enable good decisions? We argue that Bayesian methods have much to offer and use hERG-mediated QT prolongation as an example to demonstrate the advantages of Bayesian predictive/machine learning models.

Drugs that block the hERG potassium channel (KCNH2 gene) delay ventricular repolarisation, which can lead to potentially fatal cardiac arrhythmias known as Torsades de Pointes (TdP) (Shah, 2006). The duration of ventricular repolarisation is measured by the QT interval on an electrocardiogram and hERG-blockers prolong the QT interval. Pharmaceutical companies continue to invest in both experimental and analytical methods to make better predictions of clinical QT prolongation and TdP risk (Sanguinetti and Tristani-Firouzi, 2006; Gintant, 2011), and we show how Bayesian methods provide interpretable probabilistic statements of safety risk, are extremely flexible, can incorporate many sources of information, and with modern software are as easy to use as classical prediction models.

We take the perspective of an early stage discovery project planning to test a compound in the clinic and ask: based on the potency of the compound as a hERG blocker and the predicted effective plasma exposure, what is the likely outcome in a clinical QT study? First, we build a Bayesian model to predict clinical QT interval prolongation (a binary yes/no variable) using historic hERG IC_50_ values from a functional *in vitro* assay and C_max_ values from clinical studies. Next, we incorporate background knowledge and information from the literature as constraints on parameters. Finally, we use the model to predict QT risk for hypothetical new compounds and to rank a set of compounds. We show the benefits of examining uncertainties as probability distributions instead of point predictions of the best estimate.

## Methods

### Data

The data are from Pollard et al. (2017) and consist of 24 compounds. Two long-acting *β*_2_-adrenoceptor agonists were excluded from this analysis as the model is specifically designed to predict hERG-mediated QT prolongation—not all mechanisms that alter the QT interval. One might argue that a future test compound might be a *β*_2_ agonist, and by excluding these two compounds, the test compound would be incorrectly classified as low risk. However, if the test compound has a high hERG IC_50_ and low C_max_, then it should indeed be classified as low risk for *hERG-mediated QT prolongation*. It is better to tailor a model to make specific predictions instead of trying to predict another mechanism based on the data of only two compounds. A more comprehensive model could easily include assays of other ion channels, *β*_2_-adrenoceptor binding, and other mechanisms deemed relevant. Therefore only 22 of the original compounds were used, 11 of which increased the QT interval in humans.

The data consist of a binary variable indicating if a compound increased QT (defined as the upper one-sided 95% confidence interval of QT prolongation *>* 10 ms) from human thorough QT or single ascending dose studies. In addition, hERG IC_50_ values from whole-cell patch clamp experiments and C_max_ values are available. Further details about the data and compounds can be found in Pollard et al. (2017).

hERG IC_50_ and C_max_ are retained as separate variables throughout. A safety margin such as hERG IC_50_/C_max_ is not used as it assumes a constant risk for a given margin, regardless of the underlying IC_50_ and C_max_ values. For example, the following IC_50_ and C_max_ values all have a margin of 1: 1/1, 10/10, 100/100, but their QT risks could differ. A constant risk for a given margin appears reasonable with this data, but in general, this assumption is unnecessarily restrictive as two degrees of flexibility are lost when building a predictive model (another main effect and one interaction effect can be estimated when keeping the variables separate). Furthermore, human C_max_ is usually unknown when making a prediction of QT risk; only a predicted C_max_ value is available. It is therefore useful to keep C_max_ as a separate term from the experimentally measured hERG IC_50_ so that we can observe how QT risk varies with changing C_max_.

### Bayesian models

The basic idea of Bayesian inference is to update what you know—even if this knowledge is minimal—with data. In the Bayesian framework, uncertainty about any quantity is represented with a probability distribution over a space of possible values. The distribution that represents our uncertainty in a quantity before seeing the data is called the prior distribution, or just “the prior”. The prior is then updated with data to form a posterior distribution, which reflects both our prior knowledge and what the data tells us. The posterior distribution will always be narrower than the prior because data decreases uncertainty. Once we have a posterior distribution for a quantity—usually a parameter in a statistical model—we can then incorporate this uncertainty into our predictions (which is not straightforward with classic frequentist methods).

There are several advantages to using Bayesian methods and here we focus on three that are relevant to predicting drug safety. The first advantage is that all sources of uncertainty can be incorporated into the prediction. These sources include:

1. Uncertainty in the value of the outcome given the values of the parameters. Even if the parameter values were known with certainty, the outcomes are stochastic and therefore uncertain—much like knowing that a coin is fair, but being unable to predict the exact number of tosses out of ten that will land heads.
2. Uncertainty in the parameters. The parameter values are usually unknown (i.e. there is uncertainty in whether the coin is fair) and classical frequentist methods only use the single best (maximum likelihood) estimate. But greater uncertainty in the parameters should lead to greater uncertainty in the predictions, and Bayesian methods naturally incorporate parameter uncertainty.
3. Uncertainty in the value of the predictors. IC_50_ and C_max_ values are not known exactly as they are estimated from experiments. The uncertainty in the predictors should be propagated through to the predictions, but in the examples used here we assume, as in most analyses, that the uncertainty is negligible.
4. Uncertainty in the form of the model. Predictions are usually made from a single “best” model, but often other models would make different but nearly as good predictions. One might consider combining predictions from several models, known as model averaging (Hoeting *et al.*, 1999), but we do not pursue this further in this simple example as there are only two predictor variables.

A second key advantage of Bayesian methods is that external information (not contained in the data) can improve predictions. In the examples that follow, we place constraints on parameters so they can only take values in the expected direction, and also include data from a published study as priors on parameters.

The third, and perhaps most important, advantage is the interpretability of posterior distributions and summaries derived from them. Since posterior distributions contain all the relevant information about a quantity of interest, they provide an intuitive representation of a prediction and the associated uncertainty, enabling better decisions.

The basic Bayesian statistical model is

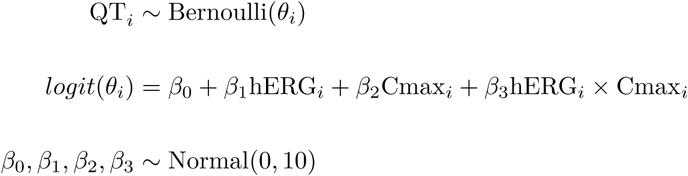

QT is the binary outcome variable indicating if a compound increased (1) or did not increase (0) the QT interval. QT is modelled as being generated from a Bernoulli distribution, and the tilde (~) is read as “is distributed as” or “is generated from” and indicates a stochastic relationship. The Bernoulli distribution is the “coin tossing” distribution, which has 1 or 0 (heads or tails) as an outcome, with the probability of obtaining a 1 given by the parameter *θ*. Thus, if *θ* = 1 we always get heads, and if *θ* = 0.5 we get heads in 50% of the cases. The subscript *i* indexes the compound and since *θ* is subscripted, this means that each compound has its own probability of increasing QT. *θ* is an unknown parameter that we want to estimate from the data, and the value of *θ* depends on the hERG IC_50_ and C_max_ values for that compound, indicated in the second line of the model definition. Since *θ* is a probability, it must lie between 0 and 1, and so the *logit* transformation maps values of *θ* to a 0-1 scale. *θ* is a deterministic function of hERG IC_50_, C_max_, and the *β* parameters (indicated with an = sign instead of a ~). The *β* parameters are also unknown and estimated from the data; *β*_0_ is the intercept, *β*_1_ quantifies the strength and direction of the relationship between QT and hERG IC_50_, *β*_2_ quantifies the strength and direction of the relationship between QT and C_max_, and *β*_3_ quantifies the interaction between hERG IC_50_ and C_max_ on QT. If *β*_3_ = 0, this implies that hERG and C_max_ have an additive effect on QT risk and that the risk is constant for a given safety margin.

In the Bayesian framework all unknowns must have a prior distribution, which represents our uncertainty about the unknown before seeing the data. The final line in the above set of equations indicates that the uncertainty in all four *β* parameters are represented as a normal distribution with a mean of zero and a standard deviation of 10.

Some compounds were tested at two doses in the clinical studies and both C_max_ values were retained in the data so that we can predict different QT risks for different exposures. Thus, the above model does not distinguish between one compound tested at two doses and two different compounds with identical hERG IC_50_ values tested at two doses.

Common metrics for quantifying the accuracy of a predictive model include the sensitivity, specificity, accuracy, and area under the receiver operator characteristic (ROC) curve, but these metrics have some limitations. The first three require an arbitrary threshold for a positive prediction, which makes inefficient use of the results from a predictive model. For example, if the threshold for a positive prediction is 0.5, then a prediction of 0.51 and 0.99 would both be classified as positive, but the strength of the prediction is lost. It is better to estimate the probability of a QT increase—a continuous value—instead of whether a QT increase will or will not happen. In addition, these metrics do not take the pre-test or baseline risk into account. For example, if 90% of the compounds are safe, then we could obtain a 90% accuracy by ignoring the hERG IC_50_ and C_max_ data and always predicting that a compound is safe. The area under the ROC curve avoids these drawbacks, but has other disadvantages (Cook, 2007; Lobo *et al.*, 2008; Hand, 2010).

An alternative metric that has not, to our knowledge, been used in the hERG literature is the Brier score (Brier, 1950), defined as

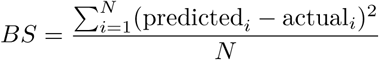

where actual_*i*_ is the 0 or 1 outcome for compound *i* and predicted_*i*_ is the value of *θ*_*i*_ from the model definition above, which lies between zero and one. The greater the difference between the actual and predicted values, the larger the Brier score, and so low scores are better. *N* is the number of compounds, and thus the Brier score is also the mean squared prediction error. Even though the accuracy would not change if a compound was correctly predicted to increase QT with a probability of 0.51 or 0.99 (assuming a 0.5 threshold), the Brier score would be much smaller with the second probability. Thus, bold predictions that are correct can be rewarded with a lower Brier score while bold incorrect predictions are penalised more heavily. Since the Brier score is calculated from *θ*, which has a posterior distribution, the Brier score also has a posterior distribution that can be used to compare the predictive ability of several models. One disadvantage of the Brier score (as well as the other methods described above) is that they do not take the number of parameters or the complexity of the model into account, and therefore more complex models will always fit the data better than simpler models. Formal methods for model comparison are available but we do not discuss them here (Vehtari *et al.*, 2016).

## Results

### Visualising the data

Figure 1A plots the safety margin (ratio of hERG IC_50_ and C_max_ values) for the 22 compounds. Some compounds were tested at multiple doses, and thus the number of points exceeds 22. Figure 1B plots the hERG IC_50_ and C_max_ values directly, and this visualisation may be preferable as information on both variables is retained and the form of the problem that we wish to solve is clear: what is the optimal boundary between the safe and unsafe compounds? Building a separating boundary on the safety margin loses up to two degrees of flexibility; the only option is to shift a vertical line left or right until an optimal classification is obtained.

**Figure 1.**
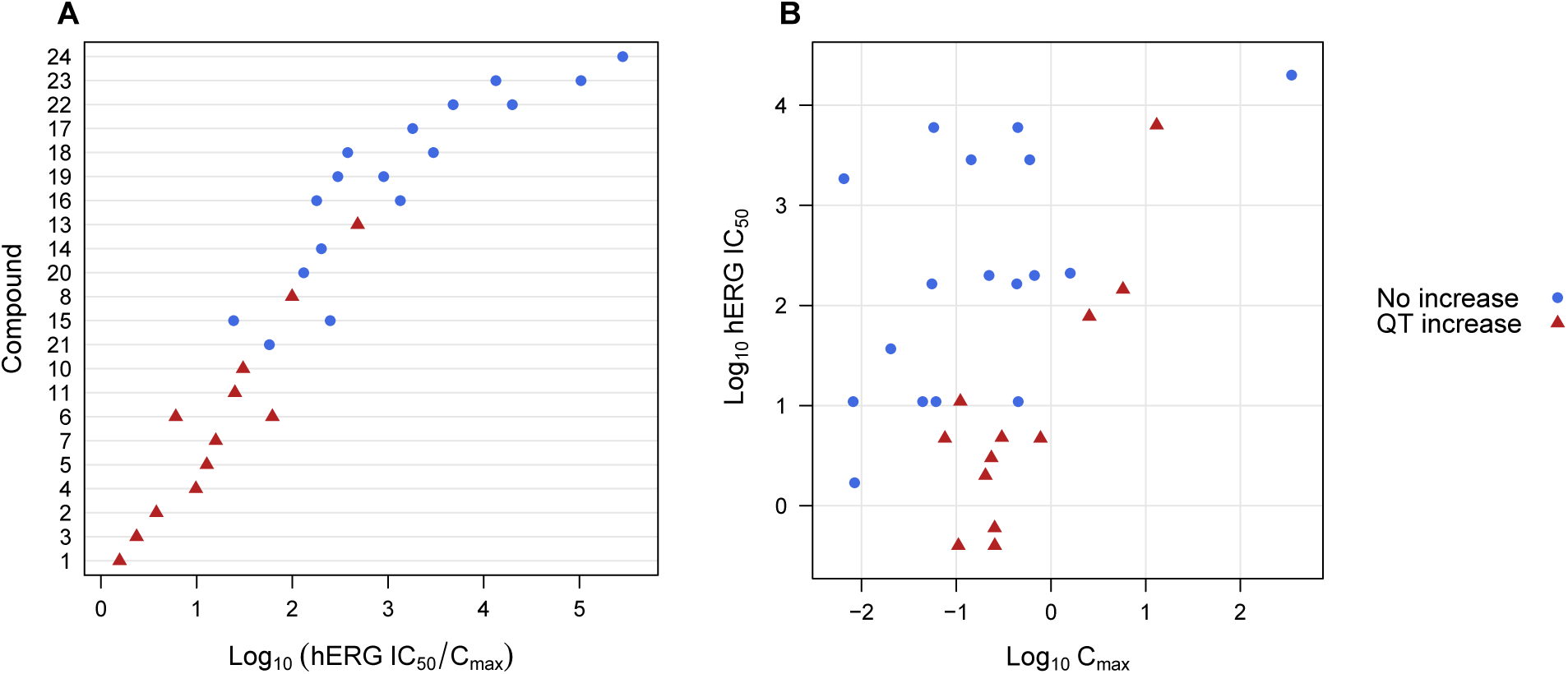
hERG IC_50_ and C_max_ values for 22 compounds. hERG IC_50_ / C_max_ margin for the 22 compounds (A), and the raw values (B). Seven compounds were tested at two doses.

The graph in Figure 1B also better highlights the relationship that QT risk increases with decreasing hERG IC_50_ and with increasing C_max_.

### Visualising uncertainty in the predicted values

From a Bayesian analysis we obtain, for each compound, a distribution of plausible values for QT risk. These predictions are shown as “violin plots” in Figure 2. Violin plots show the distribution of values, with the thickness of the distribution proportional to the density of values. These distributions are not true probability densities (their areas do not equal one) because the y-axes are compressed so that the distributions do not overlap with those above and below (see Fig. 7B and C for predictions plotted as true probability densities). Nevertheless, violin plots give an impression of where the bulk of the values lie, and many compounds can be compared. Compounds with wide distributions have insufficient evidence to make a clear prediction, which is useful to know when making a decision. Figure 2A shows the predicted probability of a QT increase when using only hERG IC_50_ values. Figure 2B includes C_max_ as another predictor, and it is clear how the distributions for most compounds shift toward either 0 or 1 because C_max_ adds information, making the predictions more certain (note: the ordering of the compounds differs in Fig 2A and B).

**Figure 2.**
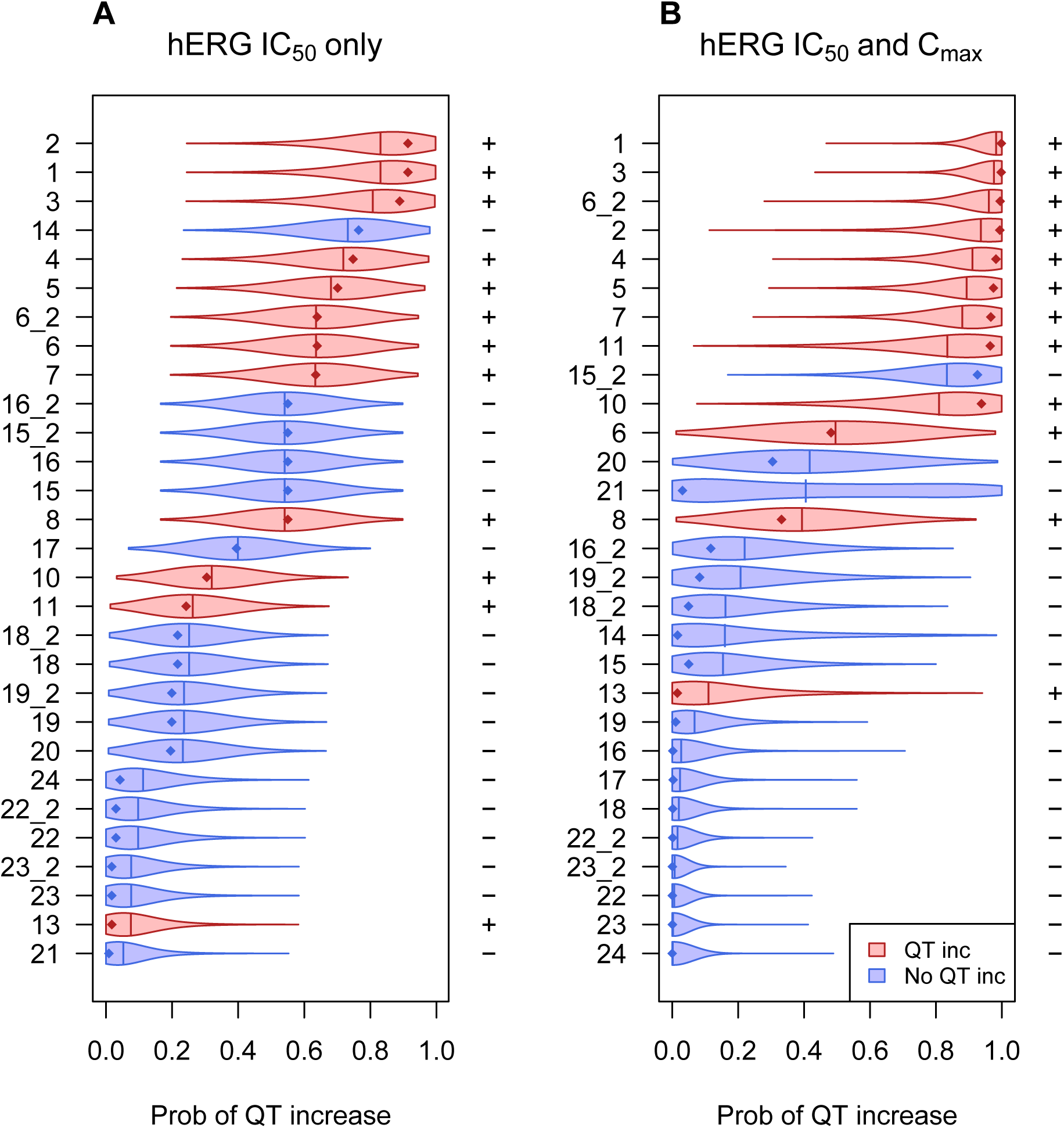
The predicted probability of a QT increase based on hERG IC_50_ alone (A) or both IC_50_ and C_max_ (B). Diamonds indicate the mode (peak) of the distribution and vertical lines indicate the mean. +/- in the right margins indicate a QT increase or no increase, respectively. Compounds are sorted by the mean of the distributions.

In Figure 2 the compounds are ranked by the mean of the distributions, but they can also be ranked by the median, mode (peak), or other summary, such as the proportion of the distribution greater than 0.5. A more compact but less informative graph could plot only summary statistics such as means and confidence intervals (usually called credible intervals to distinguish them from frequentist confidence intervals).

Figure 2B gives an impression of how the predictions improve when adding C_max_ as a predictor; compounds that increase QT (red distributions indicated with a “+” in the margin) tend to shift to the right, while safe compounds shift to the left. The Brier score captures this improved certainty in the predictions and is shown in Figure 3 for both models.

**Figure 3.**
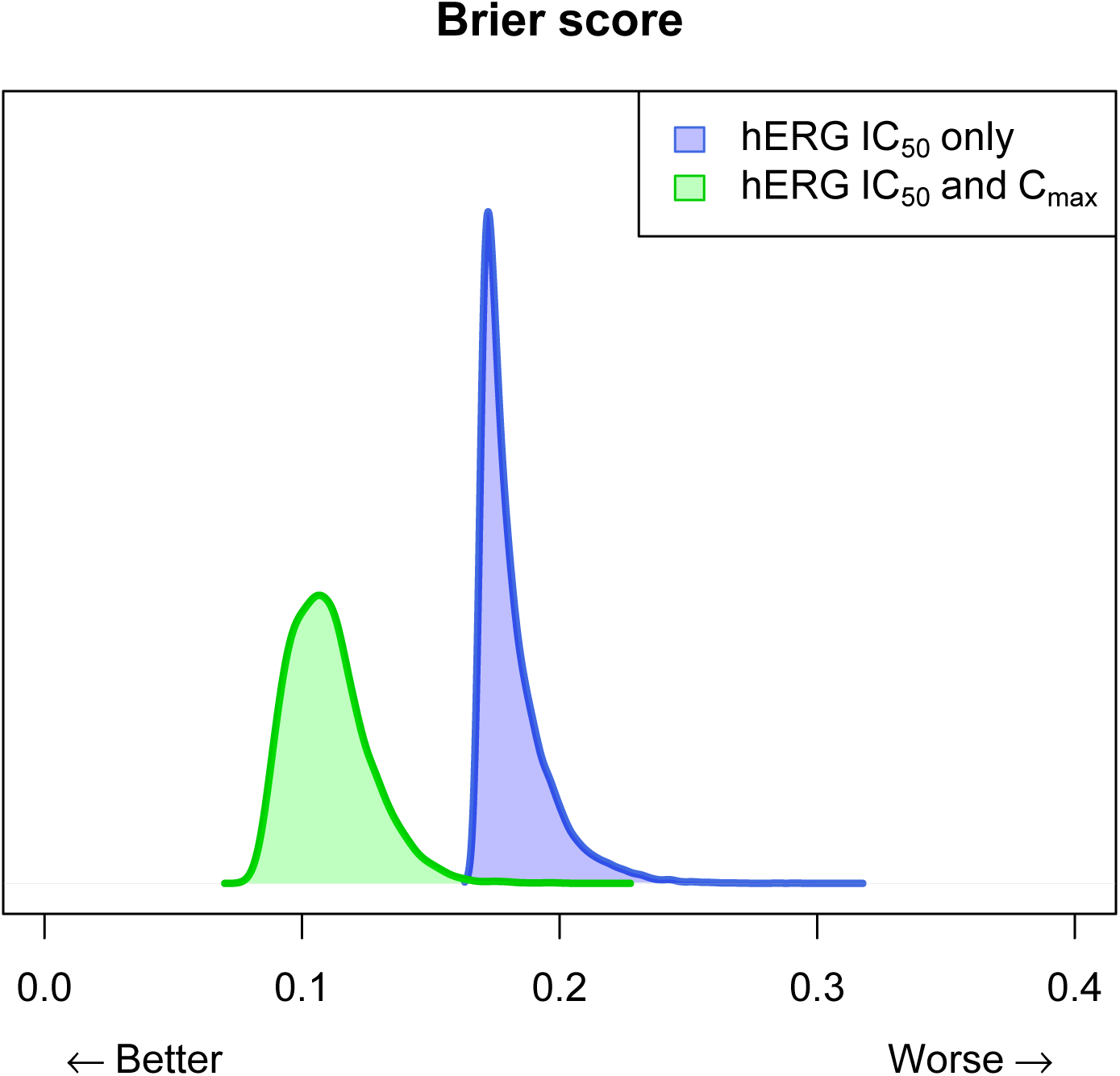
Brier score posterior distributions for a predictive model using only hERG IC_50_ values (blue) or hERG IC_50_ and C_max_ values (green).

### Incorporating background information as constraints on parameters

An advantage of Bayesian models is that prior information can be included. A simple way of including other information is by placing constraints on parameters. For example, we know that the probability of QT prolongation increases with lower hERG IC_50_ and with higher C_max_. We can state this more formally by saying *β*_1_ *<* 0 and *β*_2_ *>* 0, and we can incorporate this information into the prior distributions for these parameters. Figure 4A shows the results of draws from the posterior distribution for *β*_1_ and *β*_2_ when no constraints are applied. Although, all the points for *β*_1_ are negative, so are some points for *β*_2_, which we know should all be greater than zero. Figure 4B shows the results with the constraints, and all points fall within the desired regions.

**Figure 4.**
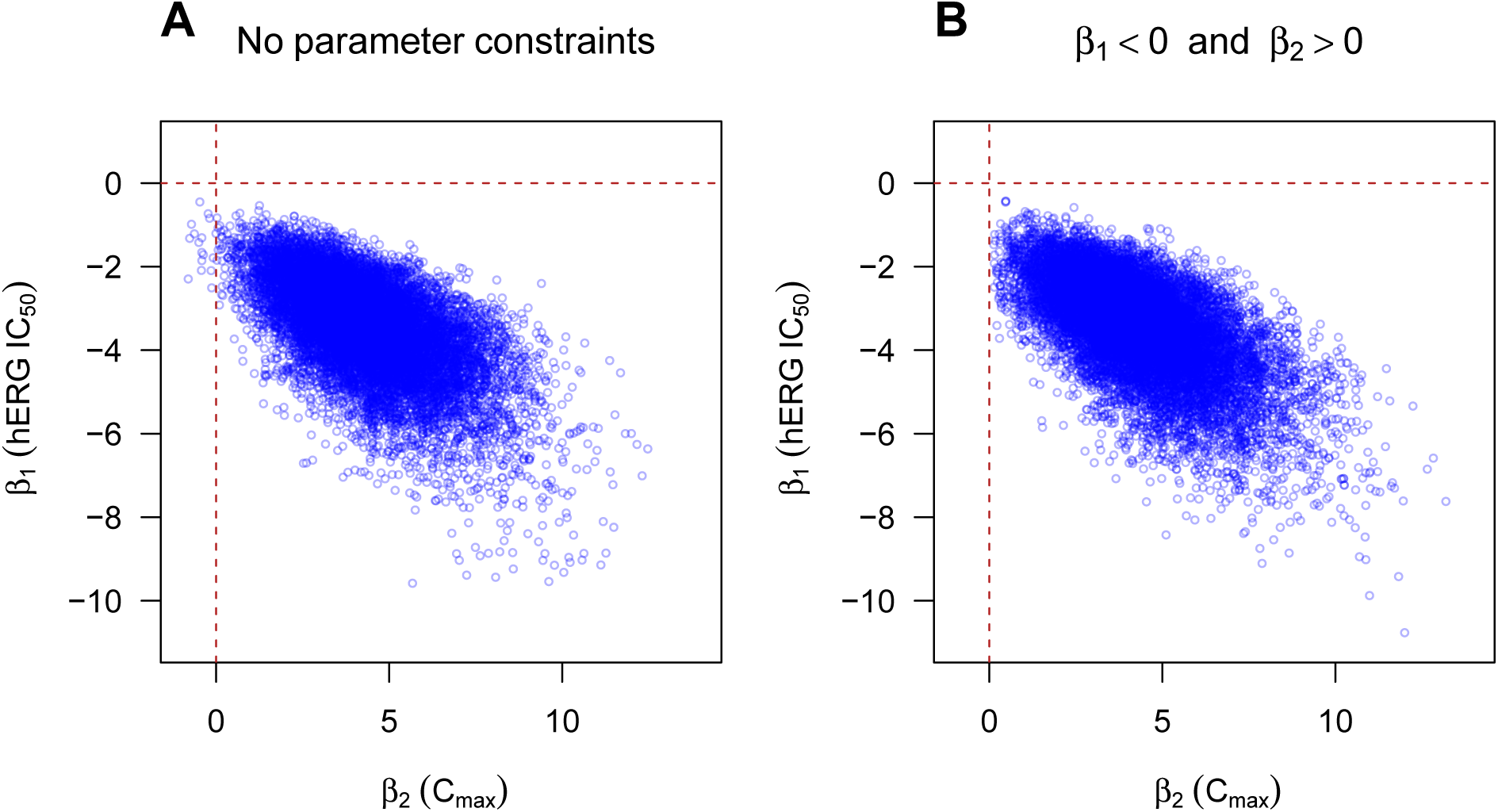
Draws from the joint posterior distribution of parameters *β*_1_ (hERG IC_50_) and *β*_2_ (C_max_). Unconstrained parameters (left) have samples for *β*_2_ below zero. Constraining parameters is trivial within the Bayesian framework and is a simple way to incorporate prior information.

Since the distribution of these parameter values are used to predict new test compounds, any change in these distributions will change the predictions. In this example the constraints make little difference, but if the relationship between the outcome and predictors was weaker or if the sample size was smaller, the uncertainty in the parameter values would be greater. This would lead to a wider spread of points, with more falling in the disallowed regions. In such a case the constraints would have a greater influence on the predictions.

### Incorporating information from the literature as prior distributions

Placing constraints on parameters is a simple way of incorporating external knowledge, but we may want to include more specific information. The extra information is from a study by Gintant (Gintant, 2011), who reported results from 39 compounds tested in humans at multiple doses. The data were manually extracted from the figures and the same model as described in the methods section was fit, but using a standard frequentist logistic model.

This additional information can be incorporated into the Bayesian model in several ways (see (Spiegelhalter *et al.*, 2004), Section 5.4) and below we describe how to include the information as prior distributions over the parameters. At one extreme, the external data may be deemed irrelevant and therefore not included. At the other extreme, the external data may be considered equal to the current data, and the easiest way of including the external data is to include it in the same data file as the current data—simply treating the external data as additional observations from the current experiment. Gintant’s data is in between these two extremes because it is highly relevant, but the physical and chemical properties of the compounds might differ between the two datasets, as might the protocols, methods, or equipment. Furthermore, as the values were manually extracted from a figure in Gintant’s paper, some additional noise was likely introduced. Thus, we treat Gintant’s data as “equal but discounted” (Spiegelhalter *et al.*, 2004) and down-weigh the information from Gintant by multiplying the standard errors of the parameter estimates by four to make them wider (this is similar to the “power prior” approach of Ibrahim and Chen (Ibrahim and Chen, 2000; Ibrahim *et al.*, 2015)). The value of four is used as an illustrative example and it reflects our opinion about how informative Gintant’s data is for the current prediction problem. The consequences of one’s assumptions and choices can be examined with a sensitivity analysis to show how the predictions change when values of the multiplier change. As the multiplier increases, the influence of Gintant’s data approaches zero, and as the multiplier approaches one, the influence of Gintant’s data is akin to treating it as extra observations from the current experiment.

After fitting a logistic model to the Gintant data, the parameters and their standard errors are: *β*_0_ = 1.59 (1.07), *β*_1_ = − 1.37 (0.51), *β*_2_ = 0.76 (0.45), and *β*_3_ = − 0.34 (0.18). These values are then used to define priors for the parameters *β*_1_, *β*_2_, and *β*_3_ for the AstraZeneca (AZ) data:

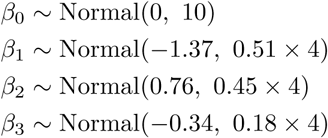

The Gintant data was not used to place a prior on the intercept (*β*_0_) as the intercept reflects the proportion of drugs in the data that have a QT risk, and there is no reason to expect this value to be similar between the datasets. We therefore used the same broad prior for the intercept as in the previous analyses. For the other parameters, normal priors were used with means and standard deviations taken from the parameters and standard errors of the Gintant analysis. The prior distributions are shown in red in Figure 5, and note how the distributions for *β*_1_ and *β*_2_ are truncated at zero, reflecting our previous constraints on these parameters.

**Figure 5.**
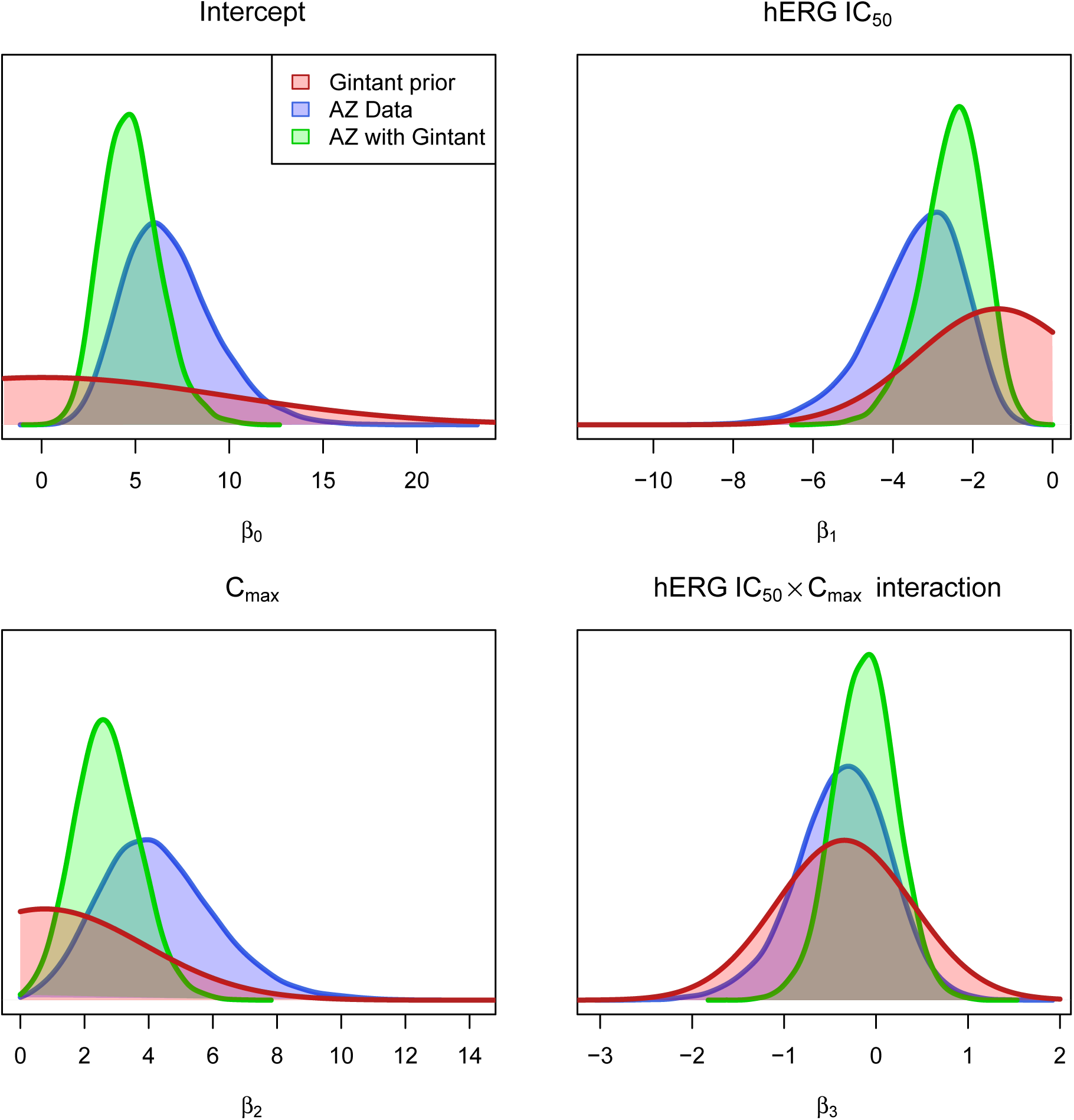
Incorporating information from the literature. Blue distributions are the posteriors with non-informative priors. Red distributions are priors for *β*_1_, *β*_2_, and *β*_3_ taken from Gintant (Gintant, 2011) but discounted (*β*_0_ used the same broad prior as the previous analysis.). Green distributions are posteriors based on priors from Gintant.

The blue distributions in Figure 5 represent the posterior distributions of AZ data without information from the Gintant study and the green distributions include Gintant’s data. The green distributions are narrower than the blue, reflecting the additional information (reduced uncertainty) that Gintant’s data provides. The distributions also have different peaks, means, medians, and so on, also reflecting the influence of external information. Since we get different distributions for the parameters when including Gintant’s data, we will also get different predictions.

The thick black line in Figure 6 is the optimal decision boundary separating safe and unsafe compounds. Since the decision boundary is calculated from the parameters (*β*_0_ to *β*_3_), which have posterior distributions, the decision boundary also has a distribution. Samples from this distribution are shown as thin grey lines, and the variability of these lines indicates the extent of other plausible boundaries.

**Figure 6.**
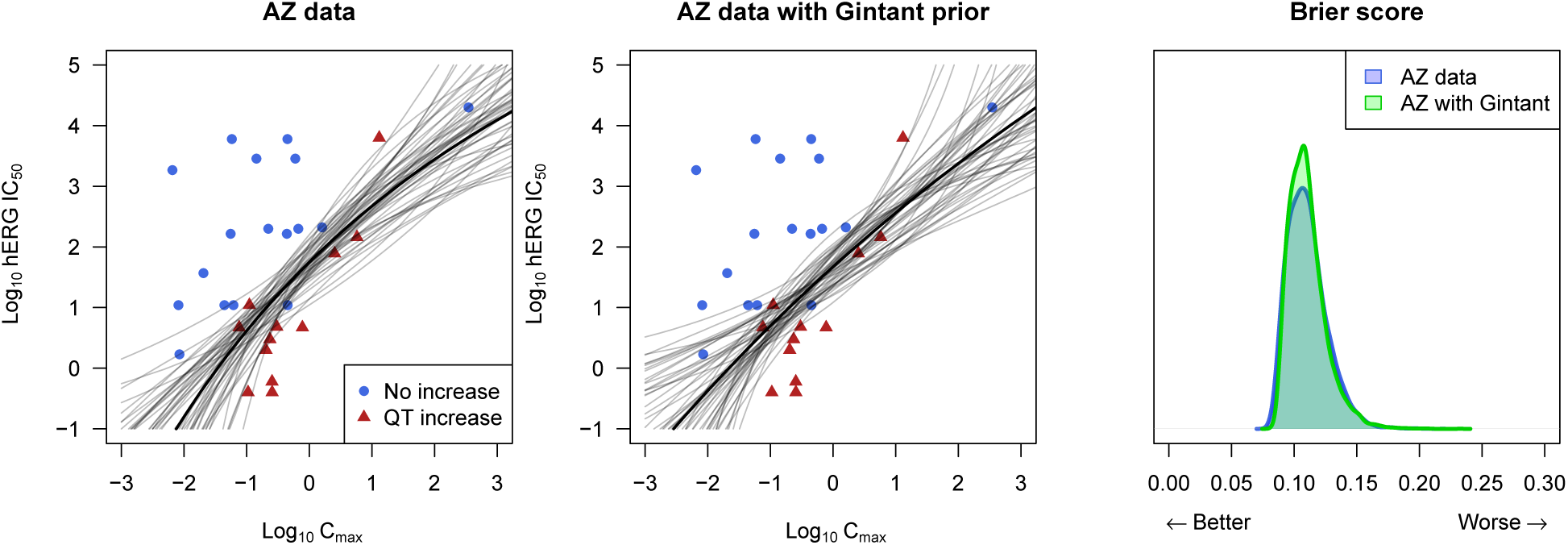
Decision boundaries and Brier scores. Grey lines are decision boundaries sampled from the posterior and indicate the extent of boundaries consistent with the data. The black line is the mean boundary. The effect of including the Gintant data is to straighten the decision boundary, which leads to one more correct classification. The distribution of Brier scores shows however that including the Gintant data has little improvement on predictions.

The middle graph in Figure 6 shows the effect of including prior information from the Gintant data set. The decision boundary changes little, but becomes less curved. As a result, one compound originally incorrectly classified as not increasing QT ends up on the other side of the decision boundary (red triangle with a Log_10_ C_max_ value of −1) thereby increasing the sensitivity and accuracy of the model. Although this may seem to be an improvement, the prediction for that compound changed little, and the Brier scores for the two models (Fig. 6, right graph) are similar.

### Predicting hERG risk for a new compound

Once a model is built from historical data, it can be used to predict the probability that a new compound will prolong the QT interval, given the compound’s hERG potency and a predicted C_max_ value. Figure 7 shows an example for a compound with a hERG IC_50_ value of 6.31 *µ*M. Since a predicted C_max_ value may be unavailable when the hERG potency data is first obtained, we plot the probability of a QT increase for a range of C_max_ values. The uncertainty in the prediction, indicated as a 90% credible interval (shaded region; Fig. 7A) is also shown. Once a predicted C_max_ value becomes available (0.02 and 0.5 *µ*M are used as an example, indicated by vertical lines in Fig. 7A), we can plot the full posteriors (Fig. 7B and C). The posterior reflects all the information we have about the probability of a QT increase, and it may be convenient to summarise the posterior with a single number, which could be the mean, median, mode (peak) of the posterior, or the proportion of the posterior that is above 50% (indicated as *P >* 0.5 in Fig. 7). For this hypothetical compound we would conclude that the risk of a QT increase is low if the C_max_ is 0.02 *µ*M and high if it is 0.5 *µ*M. We argue that such displays provide an excellent way to communicate QT risk as well as the uncertainty in this prediction.

**Figure 7.**
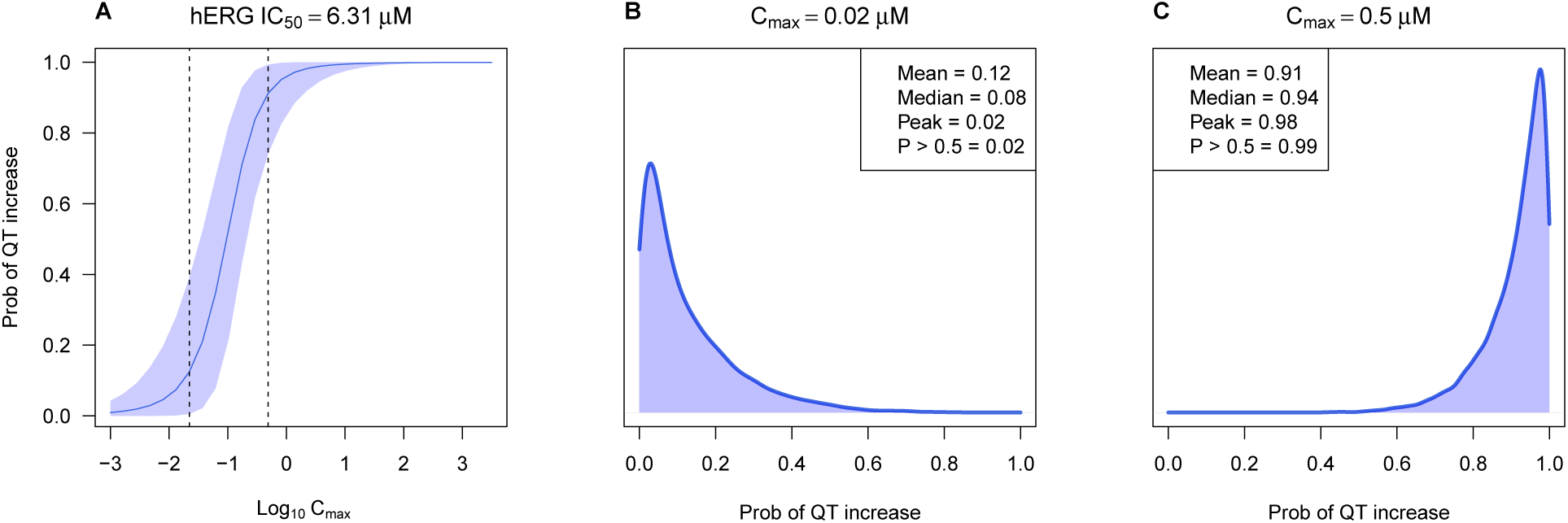
Predictions for a hypothetical new compound. If C_max_ is unknown, predictions can be displayed for a range of C_max_ values (A), and shaded regions indicate the 90% credible intervals. Alternatively, the posterior can be displayed at specific C_max_ values (B and C). Numbers in the box indicate numeric summaries of the posterior distribution.

### Ranking compounds

In drug discovery, decisions are often made to prioritise one or more compounds and exclude others. Many criteria are involved in these decisions and usually a ranked list is generated, with those at the top of the list chosen for progression. A problem with this approach is that the rankings are based on either noisy experimental measurements, predictive models with many sources of uncertainty, or both, and it is unclear how the rank ordering would change if the experiments were repeated. In other words, the stability of the ranking is unknown. Given the predictive distributions (e.g. from Fig. 2), we can easily calculate the probability that the QT risk of one compound is higher or lower than others. Figure 8A shows ten new compounds, which can either be from the same chemical series or structurally diverse, and we would like to select one as a lead compound to optimise its physical and chemical properties. The black line is the decision boundary from the original model based on only the AZ data (from Fig. 6, left). Compounds H and E are furthest from the boundary in the Northwest direction, but is it possible to distinguish between them?

**Figure 8.**
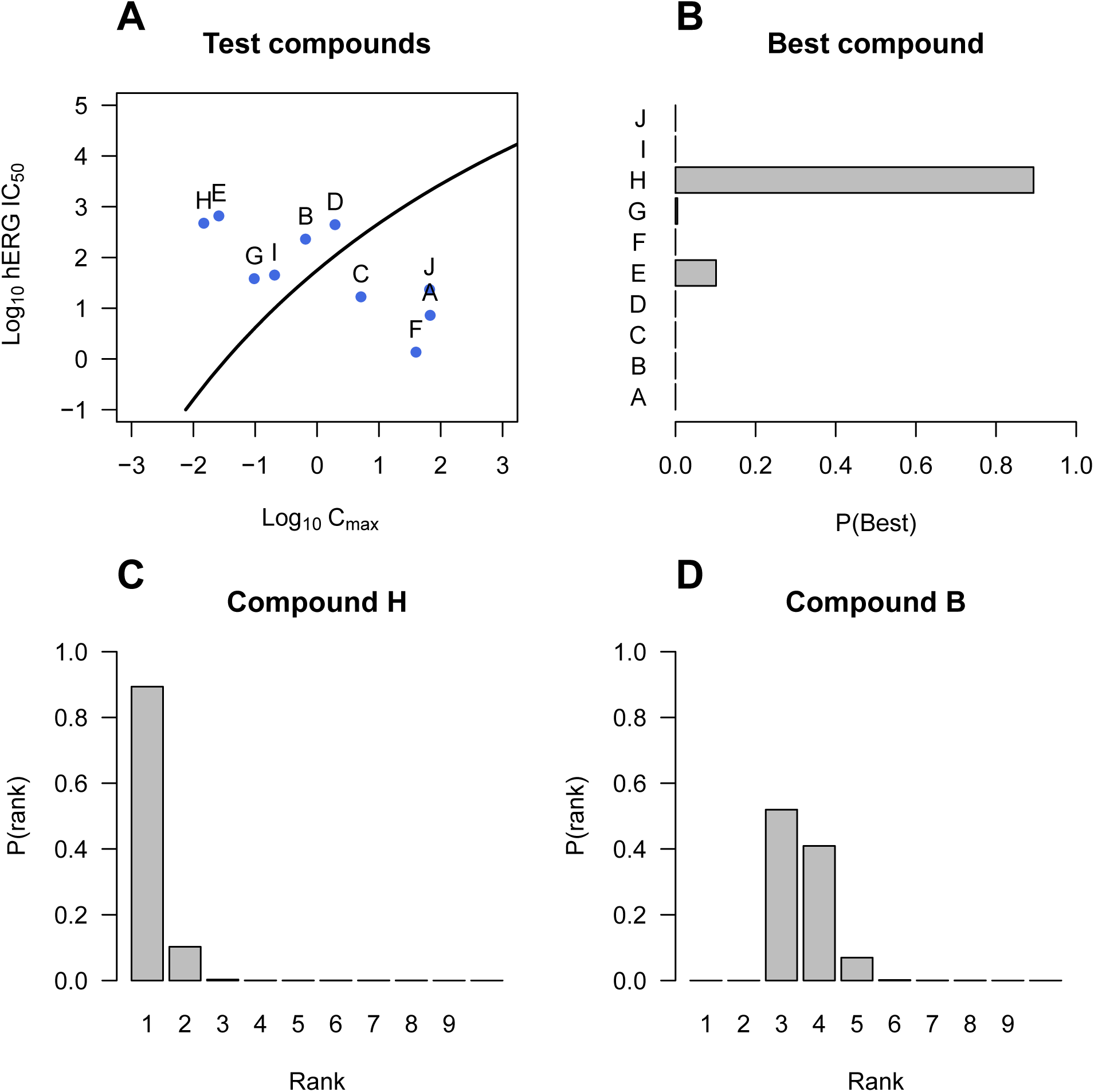
Ranking compounds. hERG IC_50_ and C_max_ values for ten hypothetical compounds and the decision boundary for QT risk (A). Probability of each compound having the lowest QT risk (B). Distribution of rankings for compound H (C) and B (D) shows the uncertainty in the rankings.

To estimate the probability that one compound is better than another, we randomly sample a value from the posterior distribution of each compound (the distributions in Fig. 2) and calculate the rankings. We repeat this process many times and calculate the proportion of times that a compound had the best rank. This is shown in Figure 8B, indicating that compound H is much better than E.

Figure 8B shows that compound H is the best but it fails to show the uncertainty in this conclusion. The distribution of rankings for compound H (Figure 8C) shows that compound H was ranked either first or second in most samples from the posterior. For comparison, the distribution of rankings for compound B is also shown, where it is less clear if it is the third or fourth best compound (Fig. 8D).

## Discussion

We used a simple example of QT prolongation with two predictor variables to highlight the benefits of a Bayesian machine learning model. Using a small historical data set, we show how a model can generate predictions of drug induced clinical hERG-mediated QT prolongation with associated uncertainty. Inclusion of this uncertainty allows risk to be contextualised and indicates how a prediction could be incorrect, enabling a framework for decision making in the non-clinical phase of drug discovery. We show how this type of data can be used to rank compounds by their predicted QT risk. Moreover, the visualisation of risk with the violin plot allows further screening paradigms to be informed. Therefore, for compounds progressing towards the clinic that are situated toward the left in Figure 2B, one might be happy to progress these through regulatory toxicology studies with reasonably high confidence that they will be devoid of hERG-mediated QT risk in the clinic. For compounds that are situated toward the right in Figure 2B, the current analysis suggests that these will be positive for hERG-mediated QT prolongation. However, several compounds have either a probability of prolonging QT of approximately 0.5, or the uncertainty of the prediction means that although the probability might be *>* 0.8 or *<* 0.2, the quality or accuracy of this prediction is too low to make a definitive statement. Under these circumstances it might be wise to supplement predictions with a more complex physiological model of hERG-mediated QT prolongation such as the anaesthetised guinea-pig (Marks *et al.*, 2012) or a large animal CV telemetry (McMahon *et al.*, 2007) to make a more accurate prediction.

Although this basic model is already useful, it can be extended in several ways to make it more powerful. First, the model can be generalised to predict overall cardiovascular risk and not just QT interval prolongation (Lester and Olbertz, 2016); for example, by using data from the Comprehensive *in vitro* Proarrhythmia Assay (CIPA) or haemodynamic/ECG data from an *in vivo* model to predict broader arrhythmia potential (Sager *et al.*, 2014). However, with more predictor variables and a fixed number of compounds (observations), there is a risk that noise will be mistaken for signal (over-fitting), leading to better predictions for the current data, but worse predictions on future data. Fortunately, fast approximations of out-of-sample prediction accuracy such as leave-one-out cross-validation and the widely applicable information criterion (WAIC) are available for these Bayesian models and can minimise over-fitting (Vehtari *et al.*, 2016). Second, uncertainty in the hERG IC_50_ and C_max_ values can be incorporated into the model (Carroll *et al.*, 2006). These values are experimentally determined and are subject to measurement error, with the size of the error likely proportional to the measured value. For example, large IC_50_ values are likely estimated with less precision because it is harder to fit a sigmoidal dose-response curve to data that do not have a clear lower asymptote. Thus, an important source of uncertainty that has traditionally been ignored can be easily included within the Bayesian framework. Third, the measured change in QT interval can be used as the outcome variable (∆QT), and not whether it was greater or less than a cut-off (upper one-sided 95% confidence interval *>* 10 ms, as used in TQT studies). Even though ICH guidelines determine the cut-off, dichotomising continuous variables loses information, can introduce bias, and does not correspond to a biologically meaningful parameter (Streiner, 2002; MacCallum *et al.*, 2002; Senn, 2003; Royston *et al.*, 2006; Fedorov *et al.*, 2009; Naggara *et al.*, 2011; Kuss, 2013). QT prolongation is a smooth function of hERG IC_50_ and C_max_ values; there is no discontinuous jump in risk between and safe and unsafe. A better approach would be to predict ∆QT directly, and then use the ICH mandated cut-off on the posterior distribution of ∆QT. As ∆QT is already collected from TQT studies, it is an inexpensive way of making the most use of the information available. Finally, QT prolongation information from non-clinical *in vivo* assessments such as the dog telemetry model could bring further information, along with a greater understanding of PKPD relationships in these more complex models.

This approach supplements current informal methods, such as looking at the hERG IC_50_/C_max_ safety margin and is straightforward to implement with modern software. For example, the first line of R code below fits the Bayesian model defined in the Methods section to the data using the `stan_glm()` function from the `rstanarm R` package (Stan Development Team, 2016b). The second line of code generates the predictions shown in Figure 7, where `new_values` are the hERG IC_50_ and C_max_ values of a new compound.

~~~
model <- stan_glm(QT ~ herg * cmax, data=QT_data, family=binomial,
prior = normal(0, 10), prior_intercept = normal(0, 10))
new.pred <- posterior_linpred(model, newdata = new_values, transform=TRUE)
~~~

For more complex models or when including additional features such as constraints on the parameters or measurement error in the predictor variables, the model definition will need to be specified in a more flexible Bayesian modelling language such as Stan (http://mc-stan.org, (Stan Development Team, 2016a)). As most safety pharmacologists are unfamiliar with R or Bayesian modelling software, at AstraZeneca we have implemented the model in a web application using Shiny (Chang *et al.*, 2017), where users input a hERG IC_50_ and C_max_ value for a new compound, and distributions like those shown in Figure 7B and C are returned. In addition, the new compound is added to a plot like Figure 1B so that it can be visually compared to compounds of known clinical risk. The data and R code are provided in the Supplementary Material to make it easier for others to implement these methods. In addition, the basic model is also coded in PyMC3 for Python users and provided in the Supplementary Material.

The current model shows the versatility of the Bayesian approach to inform decisions throughout the drug discovery process. The QT-hERG dataset provides a convenient proof of concept for these methodologies, and a similar approach can be applied to many of the decisions made in drug discovery and development, be it in the safety arena or when determining the likelihood that a molecule or mechanism might be efficacious. Crucial to every decision we make is the probability that the decision is incorrect, and transparent measures of decision accuracy enables prioritisation of compounds using all available data.

## Supplementary material

The data, R, and Python code are provided in a single zip file on Github: http://stanlazic.github.io/supplementary/ToxSci2017_data_and_code.zip

